# 3D Neuronal Mitochondrial Morphology in Axons, Dendrites, and Somata of the Aging Mouse Hippocampus

**DOI:** 10.1101/2021.02.26.433056

**Authors:** Julie Faitg, Clay Lacefield, Tracey Davey, Kathryn White, Ross Laws, Stylianos Kosmidis, Amy K Reeve, Eric R Kandel, Amy E Vincent, Martin Picard

**Affiliations:** Wellcome Trust Centre for Mitochondrial Research, Translational and Clinical Research Institute, Faculty of Medical Sciences, Newcastle University, Newcastle upon Tyne, UK; Electron Microscopy Research Services, Newcastle University, Newcastle, UK; New York State Psychiatric Institute, New York, NY 10032, USA; Zuckerman Mind Brain Behavior Institute, Kavli Institute for Brain Science, Department of Neuroscience, Howard Hughes Medical Institute, Columbia University, New York, USA; Division of Behavioral Medicine, Department of Psychiatry, Columbia University Irving Medical Center, New York, NY, USA; Department of Neurology, The Merritt Center and Columbia Translational Neuroscience Initiative, Columbia University Irving Medical Center, New York, NY, USA

**Keywords:** 3D reconstruction, mitochondria, hippocampus, morphometry, aging

## Abstract

The brain’s ability to process complex informations relies on the constant supply of energy through aerobic respiration by mitochondria. Neurons contain three anatomically distinct compartments – the soma, dendrites, and projecting axons – which have different energetic and biochemical requirements, as well as different mitochondrial morphologies in cultured systems. Here we apply a quantitative three-dimensional electron microscopy approach to map mitochondrial network morphology and complexity in the mouse brain. We examine three neuronal sub-compartments – the soma, dendrites, myelinated axons – in the dentate gyrus and CA1 of the mouse hippocampus, two subregions with distinct principal cell types and functions. We also establish compartment-specific differences in mitochondrial morphology across these cell types between young and old mice, highlighting differences in age-related morphological recalibrations. Overall, these data define the nature of the neuronal mitochondrial network in the mouse hippocampus, providing a foundation to examine the role of mitochondrial morpho-function in the aging brain.

## Introduction

The mammalian hippocampus is a specialized brain structure underlying the cerebral neocortex with a role in encoding episodic memories and spatial maps, crucial features for organizing adaptive behavior (Hartley et al., 2014). Among the four main subregions of the hippocampus, the *cornu ammonis* 1 (CA1) and the *dentate gyrus* (DG) play distinct roles in the conversion and storage of long-term memory (Kandel, 2009; Gilbert et al., 2001), and are highly specialized in both cellular and network features. For example, the DG contains many small non-pyramidal excitatory neurons, granule cells, that are much more sparsely active than the large pyramidal neurons of the CA1 layer (Diamantaki et al., 2016). However, a common property of both DG and CA1, and of brain tissue in general, is the high energy requirement to sustain normal activity.

Brain energy production, among other functions, relies heavily on oxidative phosphorylation within mitochondria, which is necessary for neurotransmitter release and for maintaining neuronal electrochemical gradients (Magistretti and Allaman, 2015). However beyond energy production, mitochondria also perform other key functions that are important for neuronal activity, such as Ca^2+^ homeostasis, fatty acid biosynthesis, Fe-S cluster formation (Youle and van der Bliek, 2012) and steroid hormone synthesis (Selvaraj et al., 2018). In neurons, mitochondria play a critical role in Ca^2+^ buffering (Lewis et al., 2018; Lin et al., 2019), which regulates synaptic transmission and plasticity. Prior to the establishment of terminally differentiated neural structures, the function and positioning of mitochondria also determine axonal and dendritic development (Courchet et al., 2013) and can influence axonal regeneration (Kann and Kovacs, 2007; Schon and Przedborski, 2011), illustrating the broad range of effects mitochondria have on neuronal behavior (Lee et al., 2018). Given these diverse roles, it is likely that mitochondrial features differ between principle cell types in DG and CA1, and may determine the relative susceptibility of these areas to age-related changes.

Depending on the tissue, cell-type, and location within the cell, mitochondria can vary in morphology from simple spheres to elongated and highly branched organelles (Yaffe, 1999; Picard et al., 2012; Leduc-Gaudet et al., 2015; Faitg et al., 2019; Vincent et al., 2019). These morphological transitions take place within seconds to minutes and directly impact mitochondrial functions (Liesa and Shirihai, 2013; Picard et al., 2013b). In fact, in humans, disease-causing primary mitochondrial defects can cause substantial changes in skeletal muscle mitochondrial ultrastructure (Vincent et al., 2016) and organelle morphology (Vincent et al., 2019). This, highlights the importance of defining the morphological characteristics of mitochondria in anatomically and functionally-distinct cell types and brain regions (DG vs CA1), as well as sub-cellular compartments.

Differences in neuronal mitochondrial morphology have previously been observed between axonal and dendritic subcompartments. In live imaging of cultured neurons, axonal mitochondria appear short and punctate; in contrast dendritic mitochondria are longer and more tubular in morphology (Chang et al., 2006; Lewis et al., 2018). Studies using serial section electron microscopy on fixed brains confirmed the same observation in the primary visual cortex of young mice (Turner et al., 2020), ground squirrels and young rat hippocampus (Popov et al., 2005). But we lack a quantitative characterization of the differences in both (i) mitochondrial size and (ii) morphology between subcompartments of functionally-defined brain areas like DG and CA1.

Moreover, aging may also influence mitochondrial morphology. Compared to young animals, older animals develop mitochondrial ultrastructural alterations and possibly a decrease in mitochondrial function in the brain, including mitochondrial respiratory control ratio (Yao et al., 2010) and antioxidant defenses (Mandal et al., 2012). Morhologically, different studies have reported that mitochondria are (i) enlarged in the aging frontal cortex (Navarro and Boveris, 2004) (ii) but smaller in the aging CA1 (Thomsen et al., 2018). Moreover, the effect of aging and disease on neuronal function within different subregions of the hippocampus appear to differ substantially (Finch, 1993, Barnes, 1994). For example, the CA1 region is one of the regions most affected in Alzheimer’s disease (Gómez-Isla et al., 1996, Braak et al., 2006), whereas with aging the DG exhibits a decline in the birth of new granule cells (i.e., neurogenesis) (Bondareff and Geinisman, 1976; Lemaire et al., 2000). However, no study as yet has examined the effect of aging on mitochondrial morphology in both the DG and CA1 of the same animals.

Therefore, we used 3D serial block-face scanning electron microscopy (SBF-SEM) (Denk and Horstmann, 2004) to quantitatively image large tissue areas and define mitochondrial morphology in the mouse hippocampus. DG and CA1 mitochondria from dendrites and somas, as well as incoming myelinated axons, were meticulously traced, reconstructed, and analyzed quantitatively, in both young and old animals. The strict lamination of these hippocampal areas represents a useful model to study compartmental differences, as it consists of distinct layers containing primarily dendrites, soma, or axons from a relatively homogeneous population of excitatory principal neurons. By doing so, we hoped to better understand cellular and network-level changes that occur in memory-related areas during aging.

## Results

### Compartment-specific imaging of neuronal mitochondria in CA1 and DG regions

Young (4 months, n=3) and old (18 months, n=4) C57BL/6J mice were transcardially perfused with fixative after brief anesthesia to preserve *in vivo* mitochondrial morphology. The dorsal hippocampus (bregma=1.64mm) was then dissected, embedded and mounted for SBF-SEM imaging of the DG and CA1 from the same brain section (Figure 1A, B). Figure S1 illustrates detailed positioning of the regions of interests (ROIs) imaged to obtain myelinated axons and dendrites (DG or CA1 molecular layers), and somata (DG granular layer, CA1 pyramidal cell layer). For each animal, all mitochondria and the cell membrane (and the nuclear envelope in the soma) from 3 axons, 3 dendrites and 2-3 somata in each hippocampal subfield were fully reconstructed in 3D with an x, y resolution of 7nm and z-resolution of 30nm (400 serial images) (Figure 1C). The dendrites from the middle of the molecular layer belong to the same cell type as the layer/somata; in contrast the axons axons identified in the DG most likely arise and project from the medial entorhinal cortex, whereas CA1 axons likely arise from the CA3, rather than from the principle cell type in the regions themselves. Below we refer to axons, dendrites and somata by their site of imaging (CA1 or DG) but the origin of axons segments is important to consider when interpreting axonal differences between the areas.

**Figure 1.**
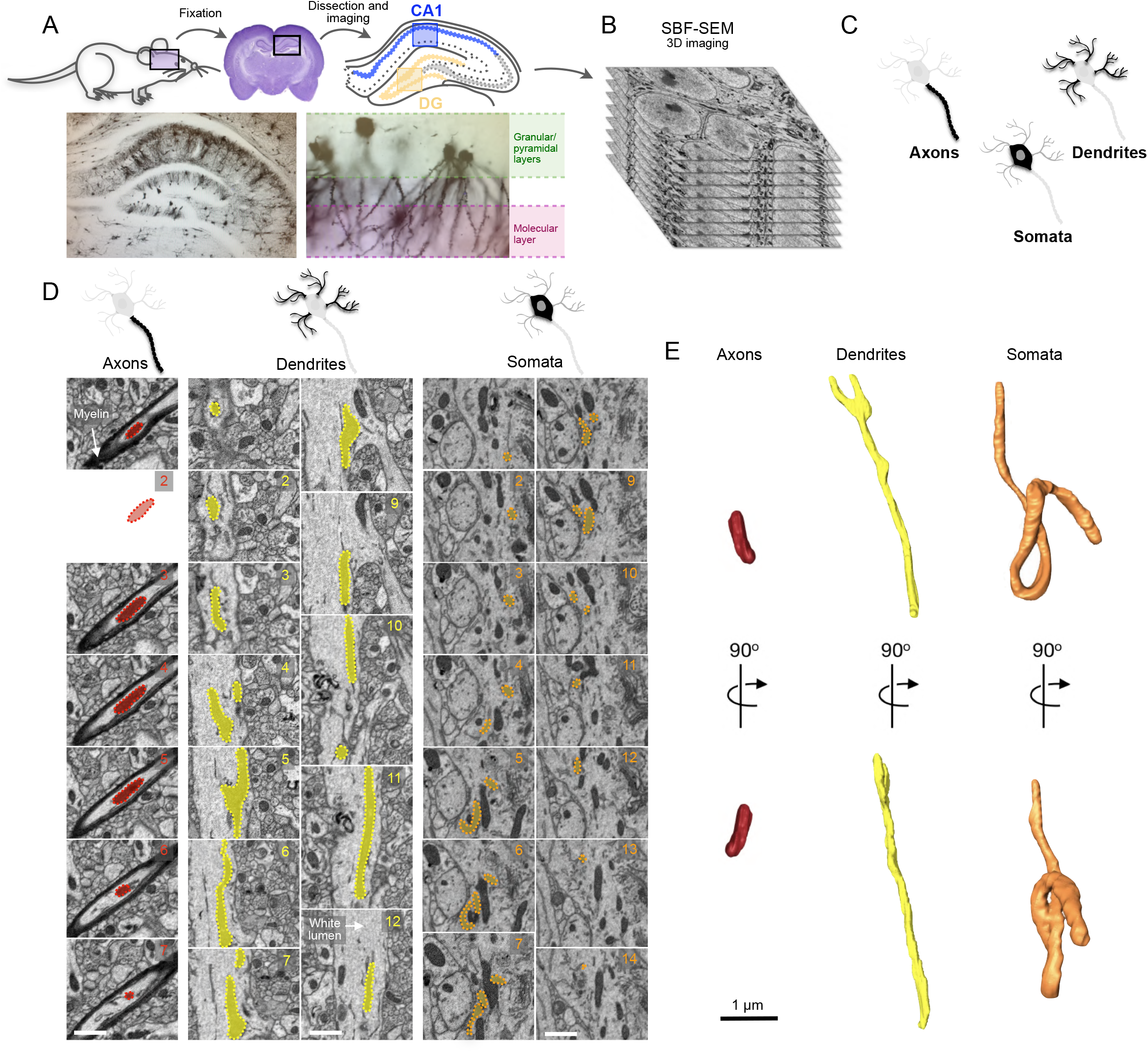
Study design for 3D reconstructions of mitochondria in neuronal sub-compartments of the mouse hippocampal DG and CA1. **(A)** Schematic of study design involving rapid transcardial perfusion and fixation, dissection, coronal sectioning, and imaging of both granular/pyramical and molecular layers in the DG and CA1. See Supplemental Figure S1 for details of sub-hippocampal regions of interest selected for imaging. **(B)** Z stack at EM resolution from serial block face scanning electron microscopy (SBF-SEM) used for 3D reconstructions. **(C)** Schematic illustrating the three neuronal sub-cellular compartments analyzed, including axons, dendrites, and somata. **(D-E)**Serial SBFSEM images sections from the CA1 numbered sequentially, showing a highlighted mitochondrion within each neuronal compartment : axons (the dark myelin on the axons), dendrites and somata **(D). (E)**Same mitochondria than in **(D)** their associated 3D reconstruction. The same mitochondria are shown seen in two different orientation (90°C rotation). Scale bars are 1 μm.

Within the molecular layer, axons were identified by the presence of surrounding myelin while dendrites were illustrated by their non-myelinated shafts with identifiable dendritic spines through the field of view. Within the granular layer, we selected for analysis only neuronal somata containing the initial segment of dendrites and axons, complete from end-to-end across the image stack so that all mitochondria within a given soma were reconstructed. In total, we analyzed 42 myelinated axons (DG: n=21; CA1: n=21), 42 dendrites (DG: n=21; CA1: n=21) and 33 somata (DG: n=19; CA1: n=14). From these compartments, a total of 1,573 (DG) and 3,892 (CA1) mitochondria were analyzed (see Table S1 for details by sub-compartment). To circumvent potential bias related to the effects of age and regional variation, reconstructions and data analysis were performed blinded to age group. For each reconstructed mitochondrion, the total volume and surface area was quantified using AMIRA, and used to compute morphological complexity (see *Materials and Methods*). Figures 1D-E illustrate typical images of reconstructed CA1 axon, dendrite, and soma mitochondria.

### Mitochondrial size and morphology across sub-cellular compartments

We first compared mitochondrial size in axons, dendrites and somata of both the DG and CA1 in young mice (Figure 2A-H). In DG, dendritic mitochondria of granule cells (Movie S1) were on average 116% larger (mean volume = 0.27 μm^3^) than axonal mitochondria in afferent axons (0.12 μm^3^, p=0.02) (Movie S2), and 42%larger than somatic mitochondria (mean volume =0.19 μm^3^, p=0.04) (Movie S3). A small proportion of dendritic (9.1%) and somatic mitochondria (3.8%) were exceptionally large (>1μm^3^), whereas no axonal mitochondria (0%) reached 1μm^3^ (Figure 2A, B), highlighting gross differences in the distribution of mitochondrial volume between sub-cellular compartments.

**Figure 2.**
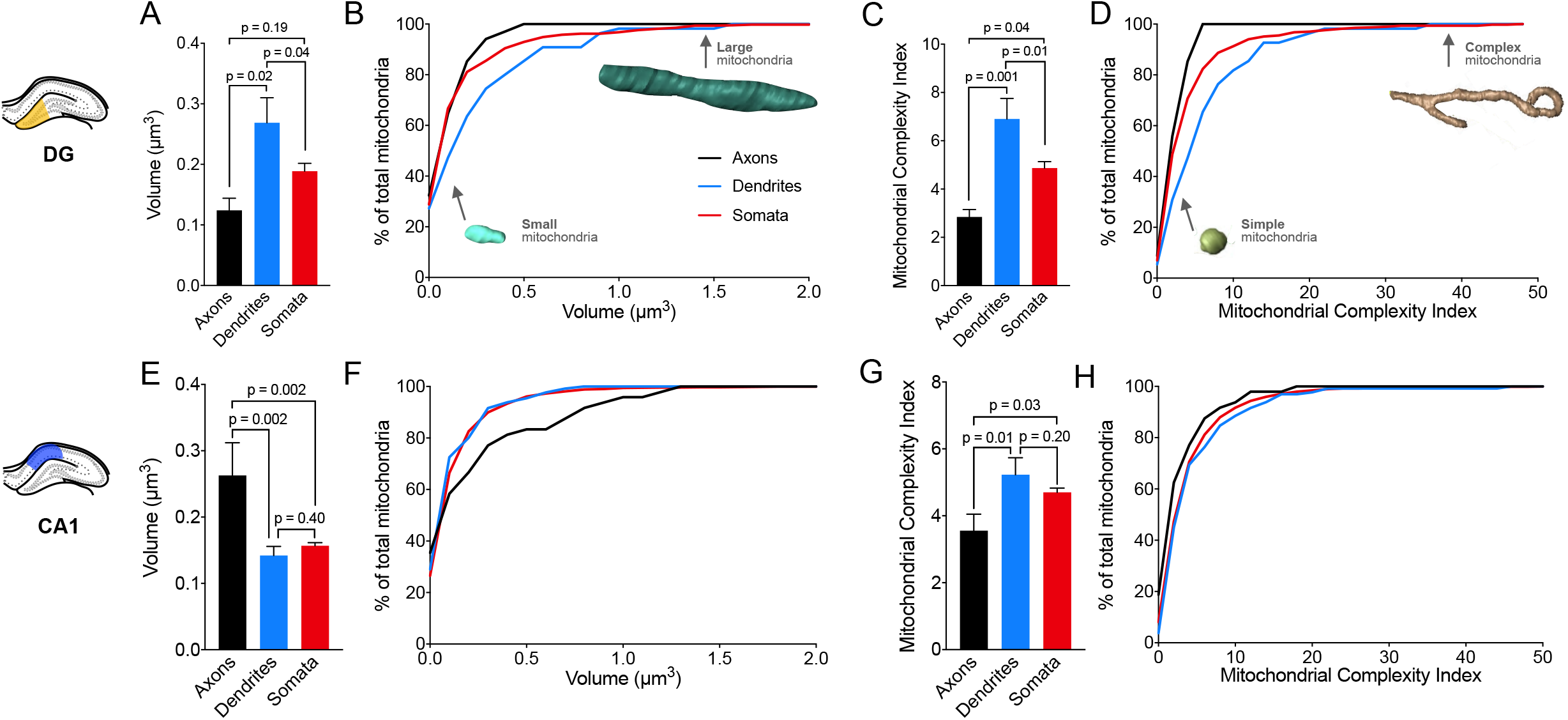
Differences in mitochondrial morphology between axons, dendrites, and somata. **(A-B)** Average DG mitochondrial volume **(A)** and cumulative frequency distribution **(B)** in axons, dendrites and somata. **(C-D)** Same as (A-B) for mitochondrial complexity index (MCI). Example mitochondria at different portions of the frequency distributions are shown for reference. n= 34 axons; n=55 in dendrites; n=471 in somata, across 3 axons, dendrites and somata. **(E-F)** Average CA1 mitochondrial volume **(E)** and cumulative frequency distribution **(F)** in axons, dendrites and somata. **(G-H)** Same as (E-F) but for MCI. n = 48 axons; n=131 in dendrites; n=1588 in somata, across 3 axons, dendrites and 2 somata. Data are means ± SEM. One–Way ANOVAs followed by post-hoc tests using the two-stage step up method of Benjamini, Krieger, and Yekutieli to correct for multiple comparisons (*p* < 0.05, *q* < 0.05).

To quantify mitochondrial morphological complexity, we next applied the mitochondrial complexity index [MCI=((SA^1.5^)/4πV))^2^] where SA is surface area and V is volume (Vincent et al., 2019). The MCI is similar to sphericity and scales with mitochondrial shape complexity, including branching and surface area relative to volume (Vincent et al., 2019). Based on this metric, in the DG, dendritic mitochondria were the most complex (mean MCI=6.9) of the three compartments, with an average MCI 143.3% greater than axonal mitochondria (mean MCI = 2.8, p=0.001), and also 41.7% greater than somatic mitochondria (mean MCI=4.8, p=0.01) (Figure 2C, D). This overall difference is mostly due to a proportion of highly complex mitochondria (MCI > 7) in dendrites (34.5%) and somata (17.6%), compared to a complete absence of complex mitochondria in axons (Figure 2D).

Interestingly, in CA1, dendritic mitochondria of CA1 pyramidal cells (Movie S4) displayed significantly lower volume (mean volume=0.14 μm^3^, p=0.002) than axonal mitochondria from afferent axons, which were on average 84.7% larger (mean volume=0.27 μm^3^) (Movie S5), and no difference was observed in the dendrites and axons compared to somatic mitochondria (mean volume=0.15 μm^3^, p=0.40) (Movie S6). Axonal mitochondria were on average 67.5% larger than somatic mitochondria (p=0.002) (Figure 2E, F). Similar to the DG, despite their larger size, axonal mitochondria were the least complex (mean MCI = 3.5) compared to either dendritic mitochondria (mean MCI = 5.2, p=0.01) or those in the somata (mean MCI = 4.7, p=0.03) (Figure 2G, H).

These data confirm previous findings that, in both CA1 and DG, axons harbor simpler mitochondria than dendrites. Moreover, mitochondrial volume and complexity in the somata of DG and CA1 regions is intermediate relative to axons and dendrites.

### DG vs CA1 differences

Following the comparison of mitochondrial morphology between sub-cellular compartments, we directly compared morphological parameters between DG and CA1, for each sub-cellular compartment.

Within the axons in the CA1, mitochondria tended to be on average 112% larger (N.S.) than in the DG, mostly driven by the presence of very large mitochondria in CA1 that are not observed in the DG (Figure 3A, B). In relation to complexity, the average MCI did not show a significant difference in axonal mitochondria between DG and CA1 (p=0.93), although CA1 axons also had 12.5% more highly complex mitochondria with MCI >7 not observed (0%) in the DG (Figure 3C, D). Dendritc mitochondria in the DG were both larger (89%, p=0.02) and more complex (MCI 31.9%, p=0.03) than in the CA1 (Figure 3G, H). For somata, both mitochondrial volume (p=0.82) and MCI (p=0.97) were similar across DG and CA1 (Figure 3I-L).

**Figure 3.**
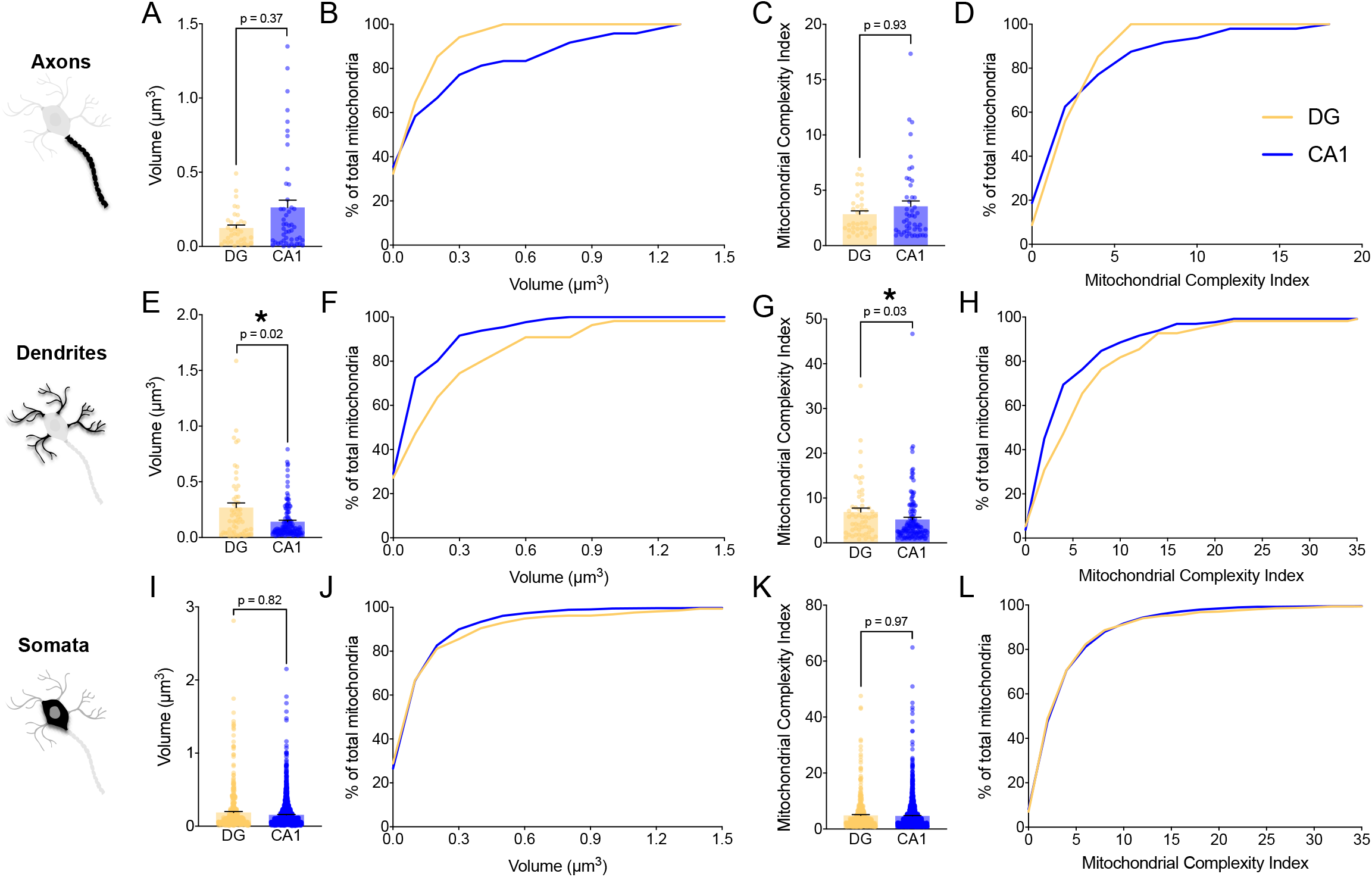
Differences in mitochondrial morphology between DG and CA1. **(A-D)** Axonal mitochondrial volume **(A-B)** and MCI **(C-D)** in DG and CA1. **(E-H)** Dendritic mitochondrial volume **(E-F)** and MCI **(G-H)** in DG and CA1. **(I-L)** Somata mitochondrial volume **(I-J)** and MCI **(K-L)** in DG and CA1. Only the dendritic mitochondria showed significant differences between the two regions, with larger and more complex mitochondria in the DG than in the CA1. DG: n= 34 axons; n=55 in dendrites; n=471 in somata, across 3 axons, dendrites and somata per mouse. CA-1: n = 48 in axons; n=131 in dendrites; n=1588 in somas, across 3 axons, dendrites and 2 somata per mouse. Data are means ± SEM. Two-tailed Mann–Whitney non-parametric tests (* *p<0.05*).

These results document the existence of notable differences in dendritic mitochondria and projecting axons between DG and CA1. By contrats, the soma of DG granule cells and CA1 pyramidal cells have mitochondria with similar morphological characteristics. This could be an important observation that reflect the function of presynaptic vs. postsynaptic dynamics in these two regions. These differences are illustrated in Figure 4A-C. In brief, axons exhibited a straight cell surface with more small simple mitochondria in DG and larger simple mitochondria in CA1 (Figure 4A). Dendrites show large complex mitochondria in DG and smaller complex mitochondria in CA1 (Figure 4B). Somata of pyramidal neurons in both CA1 and DG exhibit a population of mitochondria that are a morphological intermediary between axonal and dendritic mitochondria (Figure 4C). Plotting the mean volume and MCI together on a mitochondrial phenotype (i.e., mitotype) graph for DG (Figure 4D) and CA1 (Figure 4E) highlights the intermediate phenotype of somatic mitochondria relative to those in axonal and dendritic processes.

**Figure 4.**
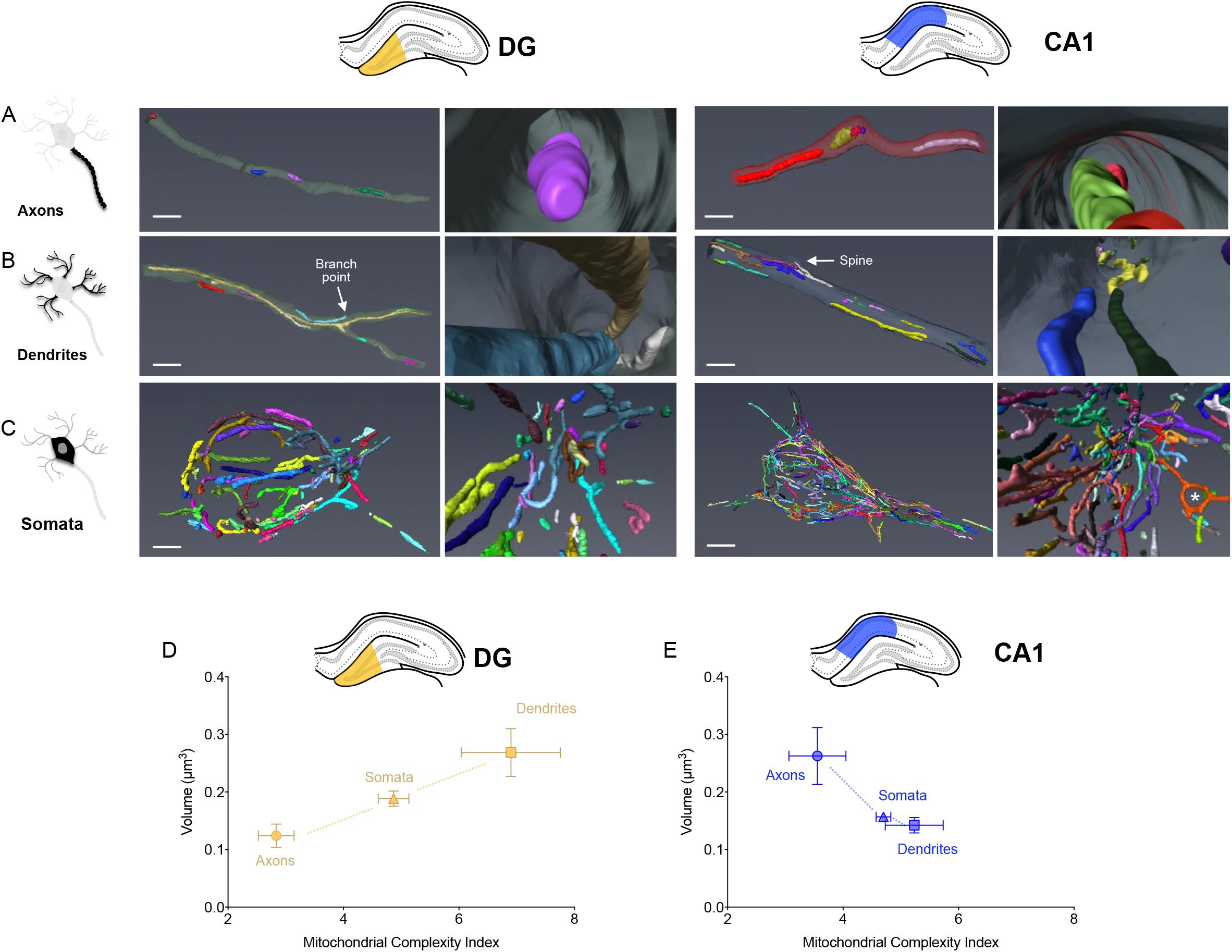
3D reconstruction of axonal, dendritic, and soma mitochondria across in the mouse hippocampus. **A-C)** Reconstruction of mitochondrial network in axons **(A),** dendrites **(B)** and somata **(C)** of the mouse DG and CA1. Each mitochondrion is a different color. Right panels are higher magnifications of the left panels. * in (C) denotes a donut mitochondrion. Scale bars are 2 μm. **(D-E)** Mitotype graphical representation of the morphological complexity (x axis) and size (y axis) of mitochondria (Mean ± SEM) in three different compartments imaged from the DG **(D)** and CA1 **(E)**.

The sub-cellular topology of mitochondria in both areas also differed between sub-cellular compartments. Whereas axonal mitochondria tended to be either distant from each other or positioned in succession along the axon lumen, in dendrites the elongated mitochondria often overlapped with each other. For example, across the diameter of single dendrites, we observed more mitochondria, particularly in proximity to dendritic spines, likely to meet regional energy demands and to contribute to local Ca^2+^ buffering (see Figure 4B in CA1). In the soma only, we noted the existence of donut mitochondria (asterisk in Figure 4C, CA1), which reflect mitochondria that loop and fuse with themselves (Long et al., 2015). As partially reflected in the MCI, somatic mitochondria harbor substantially more convoluted shapes, likely reflecting a combination of both larger available volume and more complex cytoskeletal organization relative to the filiform axons and dendrites and potentially different biochemical functions (e.g. biosynthetic).

### Differential effect of aging on axons, dendrites, and somas

To investigate the effect of mitochondrial aging in the hippocampal DG and CA1 mitochondria, we compared morphological parameters of the young mice described above to 18-month old syngenic mice. To highlight the regional differences in mitochondrial volume and complexity with aging, we used the same mitotype graph as above. The general effect of aging on mitochondrial volume and MCI is shown, separately for the DG and CA1, by the dashed arrows (Figure 5A, B, and Figure S3). As in young mice, mitochondria in axons, dendrites and somata in old animals generally had distinct mitochondrial morphologies.

**Figure 5.**
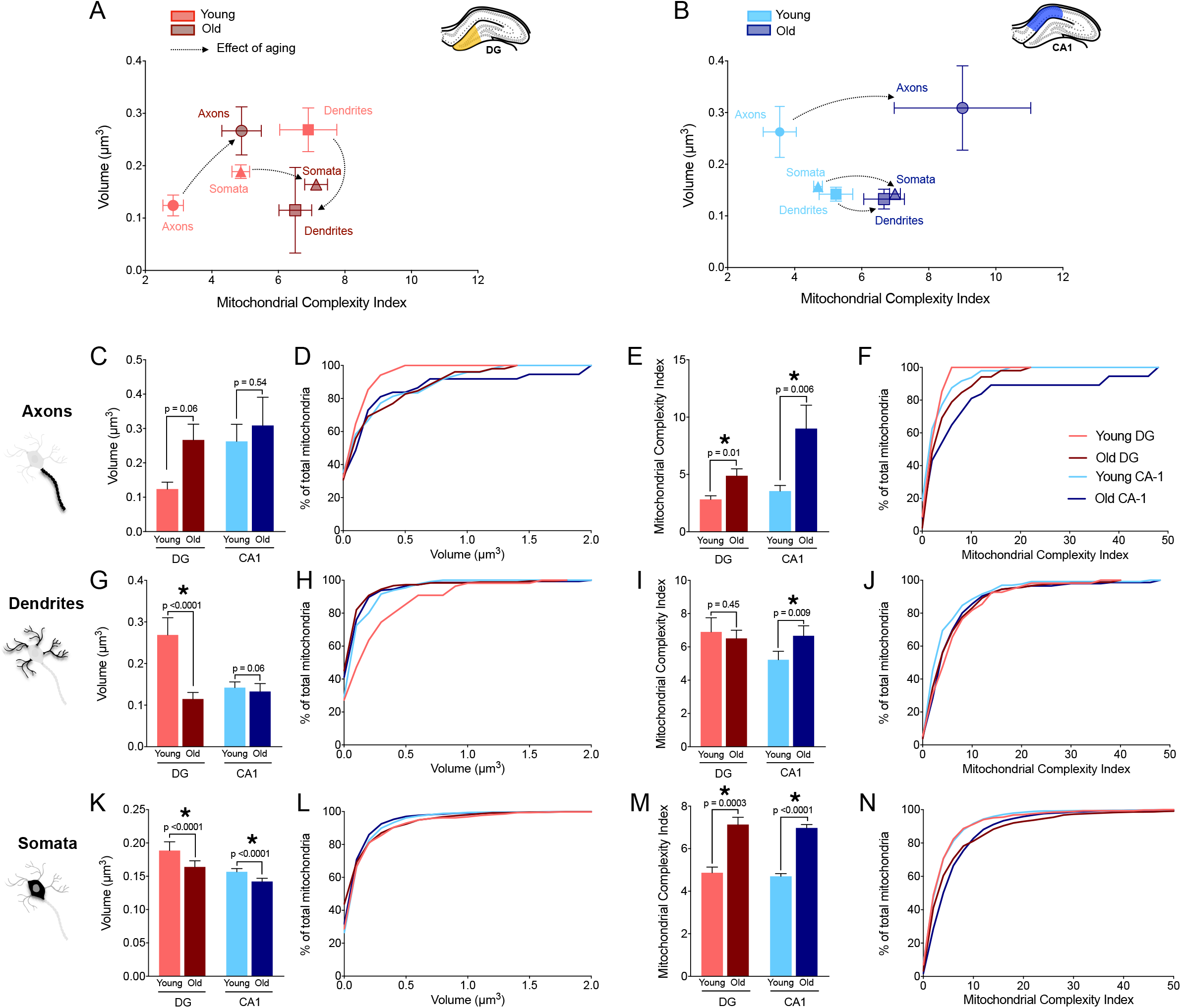
Effect of aging on mitochondrial morphology in DG and CA1 mitochondria. **(A-B)** Mitotypes illustrating the difference between aged and young mitochondria (Mean ± SEM) across compartments in the DG **(A)** and CA1 **(B)**. **(C-F)** mitochondrial volume **(C-D)** and MCI **(E-F)** comparison between young and old animals. MCI were significantly increase with age in both regions. **(G-J)** Dendritic mitochondrial volume **(G-H)** and MCI **(I-J)** comparison between young and old animals. With aging, dendritic mitochondria in the old DG were clearly smaller (mean volume=0.11 μm^3^) than mitochondria in the young DG.The MCI increased with age only in CA1. **(K-N)** Somatic mitochondrial volume **(K-L)** and MCI **(M-N)** comparison between young and old animals. With aging, the mitochondrial volume decreased significantly in both regions, however the complexity increased significantly with age. Giving smalller mitochondria but complex with age. Young DG: n= 34 axonal; n=55 dendritic; n=471 somatic mitochondria; Old DG: n= 52 axonal; n=184 dendritic; n=777 somatic. Young CA1: n= 48 axons; n=131 dendrites; n=1588 somatic. Old CA1: n= 37 axonal; n=146 dendritic; n=1930 somatic. n=3 axons, dendrites, and somata analyzed for each region (DG and CA1), in young and old (total 12 of each). Data are means ± SEM. Two-tailed Mann–Whitney non-parametric tests (* *p<0.05*).

In the DG of olf mice, contrary to young aginals, dendritic mitochondria were on average 56.8% smaller (mean volume=0.11 μm^3^) than axonal mitochondria (mean volume=0.27 μm^3^, p=0.001) and 29.9% smaller than somatic mitochondria (mean volume=0.16 μm^3^, p=0.02) (Figure S3A). Axonal mitochondria were 62.5% larger than somatic mitochondria (p=0.004) (Figure 5A, Figure S3A). Old somatic mitochondria tended to be on average the most complex, mostly driven by the presence of very complex mitochondria (13.5%, MCI>15) in somata and dendrites (9.8%), whereas fewer axonal mitochondria were very complex (5.7%) (Figure 5A, Figure S3A).

Interestingly in the CA1, dendritic mitochondria were also the smallest, on average 57% smaller (mean volume=0.13μm^3^) than in axons (mean volume=0.31, p<0.0001) and 6.9% smaller than in the soma (mean volume=0.14, n.s) (Figure 5B, Figure S3B). No differences were seen in complexity, but in contrast to young animals, the old axonal mitochondria were larger than both dendritic (35%) and somatic mitochondria (28.8%) (Figure 5B, Figure S3B).

Thus, in old animals, axonal mitochondria were the largest in both DG and CA1, but were most complex in the CA1, pointing to the differential effects of aging on both hippocampal regions. Furthermore, aging reversed the larger size of dendritic vs. axonal mitochondria in DG relative to CA1.

### Dentate gyrus mitochondria are more affected by aging than those in the CA1 region

The effect of aging on mitochondria within the DG differed between axons, dendrites, and somata. First, the average volume of axonal mitochondria did not show any significant volume differences between young and aged mice, although old animals showed a greater proportion of larger mitochondria (>0.5μm^3^, 17.3%) than did young animal (0%) (Figure 5C, D). Within the DG, old axonal mitochondria were 114.8% more complex than in young animals (p=0.01) (Figure5E, F). This was also due to a greater proportion of highly complex mitochondria (MCI > 7) in old axons (21.5%) and a complete absence of complex mitochondria in young axons (Figure 5E, F). Second, dendritic mitochondria of old animals were 58.2% smaller than in young animals (p<0.0001) (Figure5G, H). But in contrast to axons, aging did not affect dendritic mitochondrial complexity (Figure 5I, J). Third, old somatic mitochondria were 13.1% smaller (p<0.0001) (Figure 5K, L), but 46.5% more complex (p=0.0003) in the old than in the young hippocampus (Figure 5M, N).

In the CA1 region, aging did not affect the volume of axonal mitochondria (p=0.54, Figure 5C, D) but was associated with greater complexity (mean MCI=9, p=0.006), than in young animals (mean MCI=3.5, Figure 5E, F). Similarly, aging did not significantly affect dendritic mitochondrial volume (p=0.06, Figure 5G, H) but mitochondrial complexity was 27.5% higher (p=0.009, Figure 5I, J). Finally, in contrast to axonal and dendritic mitochondria, the somatic mitochondria of old animals were on average 10% smaller (mean volume=0.14μm^3^, p<0.0001) than those of young mice (mean volume=0.16μm^3^, Figure 5K, L), and also more complex (mean MCI=7.0, p<0.0001) than young CA1 somatic mitochondria (mean MCI=4.7) (Figure 5M, N).

These data indicate that aging leads to a general increase in mitochondrial morphological complexity within multiple neuronal sub-compartments from both DG and CA1, particularly incoming axons. However, compared to the CA1, DG mitochondria are in most cases more severely impacted by aging, exhibiting larger effect sizes for mitochondrial volume changes, with larger axonal mitochondria and smaller dendritic and somatic mitochondria.

### Mitochondrial volume density in aging axons and dendrites

We then broadened our analysis beyond the morphological properties of mitochondria themselves to examine their abundance (i.e., volume density) in each sub-cellular compartment of DG and CA1. Mitochondrial volume density (MVD) is calculated as the proportion of intracellular volume occupied by mitochondria, and is a direct determinant of energy production or oxidative capacity. In young brains, axonal MVD was 153.9% higher in the CA1 than the DG region (p=0.009), but the opposite was observed in dendrites where the CA1 had 45.1% lower MVD than the DG (p=0.03, Figure 6). This difference points towards potentially important differences related to DG granule and CA1 pyramidal neuron activity or biochemical needs for dendrites, and different projection sources for incoming axons.

**Figure 6.**
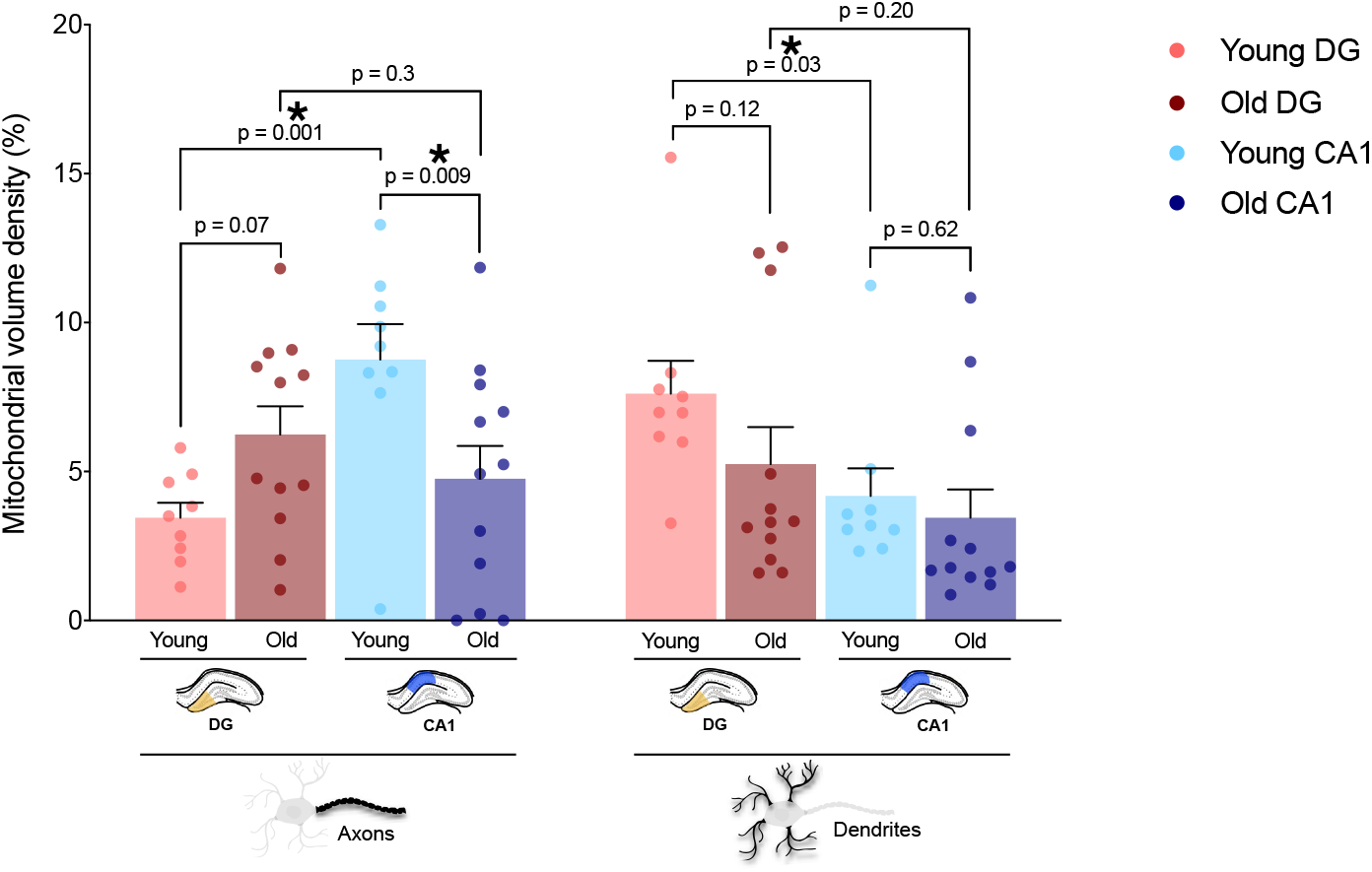
Effect of Aging on mitochondrial volume density across sub-cellular compartments. Mitochondrial volume density (MVD) in axons and dendrites in DG and CA1 for young and old mice. Young DG: n=9 axons, n=9 dendrites; Old DG: n= 12 axons; n=12 dendrites; Young CA1: n= 9 axons; n=9 dendrites; Old CA1: n= 10 axons; n=12 dendrites. Data are means ± SEM. Two–Way ANOVA followed by post-hoc tests using the two-stage step up method of Benjamini, Krieger, and Yekutieli to correct for multiple comparisons (*p* < 0.05, *q* < 0.05).

In relation to aging, MVD in old CA1 axons was 45.6% lower than in young brains (p=0.009) (Figure 6). The effect tended to be opposite in DG axons, where old DG axons had 80.9% higher MVD than young axons (p=0.07). In old dendrites of both hippocampal regions MVD tended to be 30% and 17% lower (N.S.) with aging, in DG and CA1, respectively, although we noted substantial heterogeneity and the existence of a subgroup of dendrites with remarkably high MVD (Figure 6). We next examined MDV and other cellular morphological properties specifically in the soma.

### Soma size and mitochondrial volume density differences with aging

To assess how changes in mitochondria with aging relate to overall changes in cell morphology, we quantified the total soma volume of DG granule and CA1 pyramidal neurons in young and old animals. The volume of the nucleus was also quantified to derive cytoplasmic volume (Figure 7A). In agreement with readily observed differences in the size of principal cell types in the regions, we observed marked differences in somatic volume between the DG and CA1. In the same animals, compared to the DG, the CA1 pyramidal neurons had larger nuclei (132.3%), cytoplasmic space (335.7%), and total somatic volume (220.3%) (p<0.0001) (Figure 7B). Moreover, whereas 55.6% of the cytoplasmic volume is occupied by the nucleus in the DG, this proportion is 40.3% (difference of −15.3%) in CA1 neurons (Figure 7C). These cell-level data add to the compartment-specific differences in mitochondrial morphology between CA1 and DG neurons.

**Figure 7.**
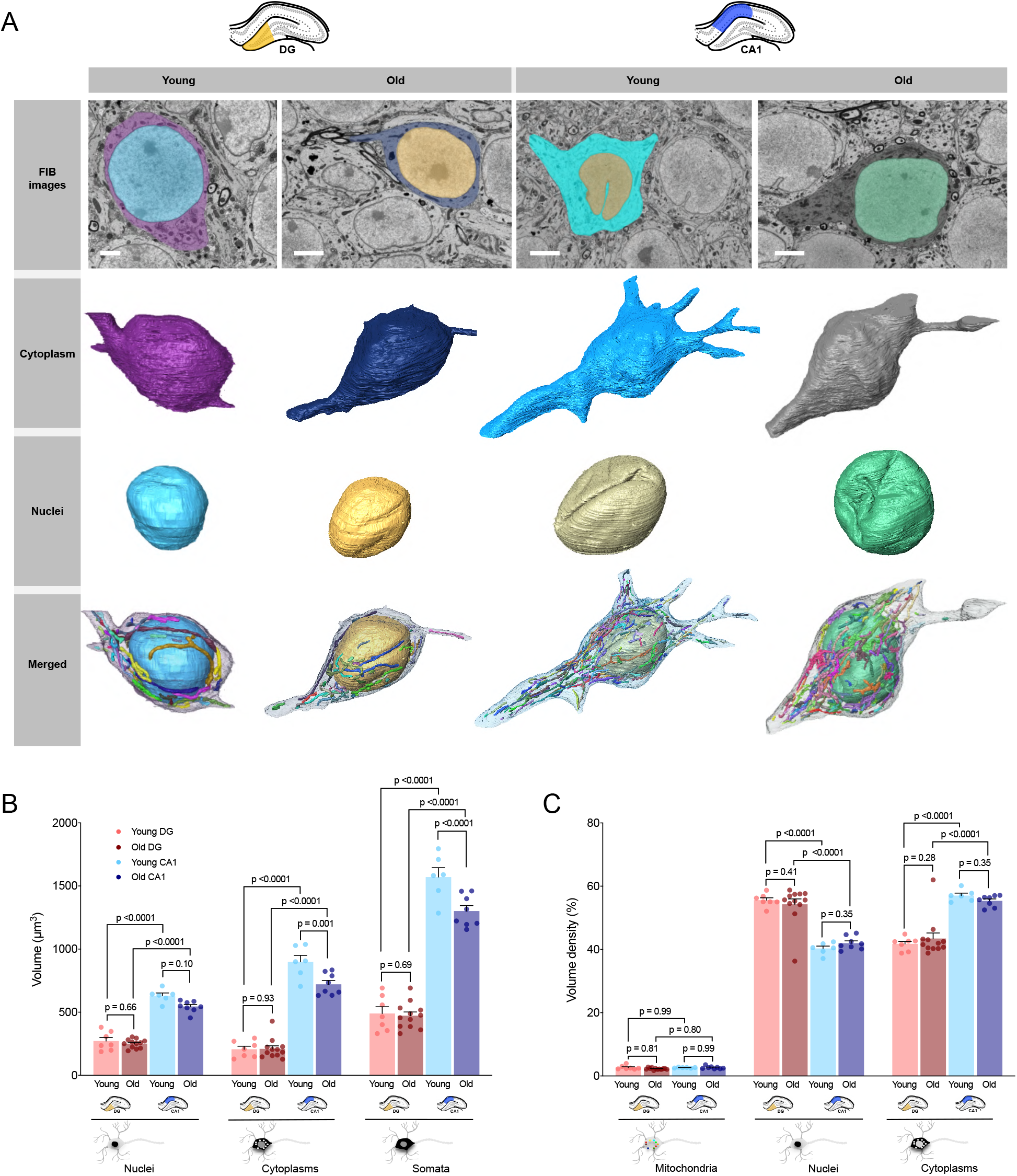
Effect of aging on sub-cellular compartments volume within the DG and CA1. **(A)** Representative 2D electron micrograph from the SBF-SEM datasets and display 3D reconstructions of the cell membrane bounding the cytoplasmic space, the nuclei, and all surrounding mitochondria. Representative images of somata from both young and old mice in the DG and CA1 are shown. Scale bars are 3 μm. **(B)** Measured volume of young and old nuclei, cytoplasms, and somata in the DG and CA1. **(C)** Volume of somatic mitochondria, nuclei and cytoplasms expressed relative to the total cellular volume, representing volume density and their differences between the young and old DG and CA1. DG: n=7 young, n=12 old cell bodies; CA1: n=6 young, n=8 old somata. Data are means ± SEM. One way ANOVA followed by post-hoc tests using the two-stage step up method of Benjamini, Krieger, and Yekutieli to correct for multiple comparisons (*p* < 0.05, *q* < 0.05).

In relation to aging, whereas DG granule neuron somatic, nuclear and cytoplasmic volume were not different between old and young animals, CA1 neurons showed atrophied soma (−17.2%, p<0.0001) and cytoplasmic volume (without nucleus and mitochondria) (−19.7%, p=0.001, Figure 7B) in old animals. Despite the apparent atrophy of CA1 somata, the volume of the nuclei was more moderately affected with aging (−13.6%, p=0.10, N.S.), indicating that old pyramidal cells have larger nuclei relative to the cell body volume.

We also computed MVD in somata, as in axons and dendrites, which showed that MVD in relatively distincg DG granule and CA1 pyramidal cells were similar. MVD also did not differ between old and young hippocampi. The percentage of total cellular volume occupied by the nucleus and cytoplasm also did not show any differences with aging (Figure 7C). Together, these data indicate that although the aging CA1 pyramidal neurons undergo substantial atrophy, the relative proportions of different cell compartment mitochondria, including MVD, are relatively unchanged in the aging mouse brain (Figure S4).

## Discussion

Using SBF-SEM 3D electron microscopy, we have quantified the distribution and heterogeneity of mitochondrial morphology in different cellular appendages of hippocampal DG and CA1 regions in young and aged mice. Using the strictly laminated structure of these regions, we were able to compare fine-grained mitochondrial features within specific subcellular compartments in these regions during aging. Our analyses confirm the structural specificity between axonal, dendritic, and somatic mitochondria and provide high-resolution quantitative information about both mitochondrial volume and morphological complexity across both DG and CA1 regions. Moreover, we find age-related recalibrations in mitochondrial morphology and soma characteristics, which both differ by subcellular compartment, and between DG and CA1 regions.

### Mitochondrial morphological specificity within neuronal sub-compartments

In neurons, mitochondria are transported to sites with high energy demand or specific biochemical requirements, such as growth cones, pre- and post-synaptic elements, and nodes of Ranvier (Hollenbeck and Saxton, 2005). Mitochondrial morphology and function (i.e., morphofunction) are interlinked in mammalian cells (Bulthuis et al., 2019; Picard et al., 2013b), and modification of mitochondrial morphology impacts neuronal mitochondrial Ca^2+^uptake and synaptic transmission (Lewis et al., 2018), positioning mitochondria as neuromodulators.

Our data confirm that mitochondria in the mouse brain exhibit distinct morphology in axons and dendrites, and adds quantitative information about these differences. Extending previous reports (Popov et al., 2005), we find that across both young CA1 and DG, axonal mitochondria were simple compared to the 65% (CA1) or 240% (DG) more complex dendritic mitochondria. This simple morphology has been shown to be important for axonal mitochondrial trafficking (Li et al., 2004), as punctate mitochondria are more easily transported along microtubules through the narrow axon lumen than tubular organelles. The function of mitochondria has also been found to differ between axonal and dendritic mitochondria, with dendritic mitochondria maintaining a greater membrane potential than axonal mitochondria (JC-1 labelling in cultured hippocampal neurons) (Overly et al., 1996). This would be in keeping with the notion that elongated, fused mitochondria may have greater coupling efficiency (Gomes et al., 2011). We also find that somatic mitochondrial volume is intermediate relative to the other neuronal compartments in both DG and CA1. In the DG, because larger mitochondria in dendrites likely arise from smaller migrating somatic mitochondria, this implicates the mitochondrial fusion machinery in the elongation of incoming organelles, possibly in addition to local biogenesis and expansion of mitochondria within the dendrites themselves (Kuzniewska et al., 2020). Furthermore, reduction of dendritic mitochondrial volume in the aged DG may be a product of defects in these processes, which could contribute to age-related memory decline.

The morphological specialization of mitochondria between different compartments of a given cell type is not unique to neurons. In skeletal muscle, subsarcolemmal and inter-myofibrillar mitochondria in the same cell also have different morphologies. In myofibers, the magnitude of morphological differences is a ~1-1.5-fold in humans (Vincent et al., 2019), and ≥1-fold in the mouse soleus (IMF mitochondria are twice as complex as SS) (Picard et al., 2013a). In comparison, in neurons we find differences of 1-2-fold between axons and dendrites, indicating that morphological specialization may be particularly pronounced in neurons. Such heterogeneity between sub-cellular compartments is in contrast with other cell types, such as hepatocytes or adrenal gland secretory cells where most mitochondria are highly similar, and where cell morphology is relatively simple consisting mostly of a uniform ovoid cytoplasmic space (unpublished data). Thus, we speculate that cell-level complexity, such as the highly specialized axons and dendrites of neurons, or the sub-sarcolemmal and intermyofibrillar compartments delimited by the contractile apparatus in muscle, may contribute to generate biochemically and energetically distinct cellular compartments, which in turn drive the establishment of specialized size, morphology, and function among resident mitochondria.

### Sub-cellular compartment influences mitochondrial morphology more than the region where it is located

It is well established that each hippocampal region possesses its unique anatomical and connectivity patterns that characterizes the region. Indeed, by finding differential distinct gene and protein expression patterns in mice, (Farris et al., 2019) identified specific mRNAs localized in the specific hippocampal subregions (CA1, CA2, CA3, and DG). These differences in transcriptome were suggested to be related to regulation of neuronal plasticity and mitochondrial function (Farris et al., 2019).

As mitochondrial morphology and function are interlinked, the divergence of mitochondrial morphology between subregions could also be due to the diverse expression of certain proteins, such as calcium binding-proteins. One of the most commonly investigated calcium binding-proteins is calbindin-D28k, known to be essential for neural function (Simons, 1988). Interestingly calbindin-D28K is more highly expressed in the DG compared to CA1 indicative of a higher calcium buffering capacity in the DG (Frantz and Tobin, 1994), which could contribute to the differences between dendritic mitochondria in DG and CA1.

As previously demonstrated by Lewis et al. (2018), a higher uptake of calcium by mitochondria is associated with reduced mitochondrial fission and elongated mitochondrial morphology. Within the cytoplasm, larger mitochondria can also take up and buffer larger amount of Ca^2+^ (Paltauf-Doburzynska et al., 2004). Here we find that dendritic mitochondria in the DG are both larger and more complex than dendritic mitochondria in the CA1, which may mean that they have a greater calcium uptake capacity and are more metabolically active than smaller axonal mitochondria, and those in CA1.

### Aging is associated with selective changes in mitochondrial morphology in the DG

Our data shows that different hippocampal regions exhibit different degrees of age-related changes. Specifically, previous work has shown that the DG may be preferentially impacted relative to the CA1 due to the capacity of the DG region for adult neurogenesis, an ongoing process that impacts DG function but declines with age (Pawluski et al., 2009). Mice exhibit age-related changes in both DG function and cerebral blood volume in the DG (Small et al., 2002), which may relate to a decline in neurogenesis. A clear finding from a proteomic study on rats examining aging related gene expression changes in the DG and CA regions also showed robust age-related changes in transcription in the DG (Ianov et al., 2017). Furthermore, the changes in the DG have been linked to age-related decline in memory (Kandel, 2009). One possible reason for this may be a decline of the histone-binding protein RbAp48, necessary in the DG for proper hippocampal function, which undergoes selective age-related deficiency in the DG in mouse and human brains (Pavlopoulos et al., 2013).

The functional impairment of mitochondrial respiratory chain activity has previously been observed in the aged brain, and may underly impairments in DG neurogenesis (Navarro et al., 2011; Holper et al., 2019). For example, ablation of the mitochondrial transcription factor A (Tfam) in quiescent neural stem cells impaired not only mitochondrial function but also neurogenic potential (Beckervordersandforth et al., 2017). Furthermore, Braidy et al. (2014) found reduced activities of mitochondrial complex I-IV in aged rodent brains. These mitochondrial impairments are the direct consequence of a decrease in the rate of electron transfer (Navarro and Boveris, 2004).

Based on the age-related decline in DG function compared to the CA1, we initially hypothesized that mitochondrial morphology may also be preferentially altered by aging in the DG compared with CA1. We find that changes in mitochondrial morphology occur in both areas, but that they were indeed more pronounced in the DG. In the CA1, mitochondrial complexity increased with age in all sub-cellular compartments. Within the DG, the MCI also generally increased with age, with one exception (dendritic mitochondria). Interestingly, DG granule cells have large dendritic mitochondria relative to DG somata/MEC axons, and CA1. This difference seems to vanish with aging. This could be a hallmark of progressive storage of information/specialization of DG neurons, and age-related defects may be a result of depleting the reservoir of dendritic/mitochondria plasticity.

It is known that mitochondrial fusion and an increase of MCI are linked with reduced mitophagy, low levels of mitochondrial dysfunction and increased calcium uptake capacity (Bordi et al., 2017; Maltecca et al., 2012; Rambold and Lippincott-Schwartz, 2011). Here, the general increase of MCI in both regions could act as a compensatory mechanism to reduce the detrimental consequences of accumulated defects that ultimately lead to respiratory deficiency. A similar phenotype of increased mitochondrial complexity with aging was also observed in aging mouse skeletal muscle (Leduc-Gaudet et al., 2015), postulated to reflect compensatory hyperfusion response to stress, as previously reported in cellular systems (Shutt and McBride, 2013). This could indicate a link between a greater age-related decline in mitochondrial function and neuronal function in the DG, which shows a large degree of age-related dendritic mitochondrial fragmentation.

### Aging preferentially impacts mitochondrial volume density in CA1 axonal mitochondria

In our study, the general lack of difference on DG and CA1 MVD between old and young animals is consistent with previous studies which showed that aging has no effect on the neuronal mitochondrial content in rats (Bertoni-Freddari et al., 1993). One exception is the axonal mitochondria in the CA1 which mostly arise from the CA3, and show a significant decrease in aged mice. While mitochondrial density did not change dramatically with age, mitochondrial morphology changed more substantially towards more complex mitochondria. Again, these changes in morphology could reflect compartment-specific and cell-type specific bioenergetic changes that relate to neuronal function in these regions. Future functional studies with both spatial and molecular specificity will be required to address these questions.

In young animals, we noted a clear difference between sub-cellular compartments, axons and dendrites across DG and CA1. Indeed, axons arising from CA3 have a higher MVD than those arising from the medio-entorhinal cortex. Also within the dendrites, the MVD in DG is higher than in CA1 dendrites. This difference points towards potentially important differences related to DG granule and CA1 pyramidal neuron activity or biochemical needs for dendrites, and different projection sources for incoming axons.

### The soma as a center for mitochondrial homeostasis?

It is currently believed that the soma acts as a central mitochondrial hub between axons and dendrites and it was demonstrated to be the site of mitochondrial biogenesis (Davis and Clayton, 1996), with mitochondria subsequently shuttled to distal axons and dendrites (Misgeld and Schwarz, 2017). At the other end of the mitochondrial lifecycle, mitophagy in neurons has also been shown to occur in the soma in Purkinje neurons (McWilliams et al., 2016), and that mitophagy of synaptic mitochondria is outsourced in a transcellular manner to surrounding glial cells (Davis et al., 2014), suggesting that mitophagy may be limited in dendrites and axons away from the soma. Here we show that the morphology of mitochondria within the soma is highly variable and that the somatic mitochondria on average present a morphological intermediate between the dendritic and axonal mitochondria. Similar findings to these were recently reported in the mouse visual cortex (Turner et al., 2020). The intermediate characteristics of somatic mitochondria suggest that they may be the source that “feeds” mitochondria to both axons or dendrites.

In our study, within the DG, mitochondria are more packed due to the smaller size of DG granule cells compared to the large pyramidal neurons in the CA1 region, which also have larger nuclei. While DG granule neurons may be smaller in terms of overall volume, and as expected the occupancy of their nucleus is higher compared to the CA1 region. The physiological significance of these findings is not immediately clear but could impact bioenergetic or electrophysiological characteristics of these cells.

With normal aging, it is now known that there is no significant loss of neurons in CA regions and low deterioration in the DG region in humans (West et al., 1994) and rodents (granule layer, hilus, CA1-2-3 and subiculum) (Rasmussen et al., 1996). The effect of aging is characterized essentially by structural and functional modifications, without overt cell loss. (Chang et al., 2004). Indeed, such defects qualify as potential morphological correlates of senescent decline in relational memory in that they can be expected to compromise the functional integrity of a region of the brain known to be intimately involved in this type of memory. Indeed, older brains tend to shrink, yet retain their neurons, which increases neuronal density (neurons per mm^3^). Whether these changes in the hippocampus are adaptive or maladaptive is unclear, but previous evidence suggests that they may represent a compensatory response to age-related changes in activity (Barnes, 1994).

In our study, the effect of aging differed both across subcellular compartment as well as by CA1 and DG subregion. With marked signs of aging in mitochondria within the DG in all subcompartments, such as fragmentation and increase in complexity which could indicate general bioenergetic dysfunction or some other process, we speculate that these features may contribute to age-related cellular and network problems in the DG that underlie memory disorders.

## Conclusion

While it is accepted that mitochondria functionally and morphologically specialize to meet the metabolic and biochemical demands of their local environment, the morphological characteristics of mitochondria in different neuronal appendages of the aging DG and CA1 had not been defined. With our study, we have confirmed that mitochondrial morphology differs between sub-cellular location, and that axonal and dendritic mitochondria are generally morphological opposites in young healthy animals. This could be related to different roles of mitochondria in either energy supply and/or calcium buffering between sub-compartments. Quantification of mitochondrial morphology in young and old animals brains showed particularly marked differences in the DG, consistent with previous evidence indicating that the DG is more pathologically affected with aging. The CA1 specifically showed atrophy of neuronal somata, pointing to diverging cellular and metabolic recalibrations between DG and CA1 neurons. Together, our findings lay the foundation to investigate potential mechanistic links between mitochondrial morphology and neuronal function, together with circuit-level function and related behaviors.

## Acknowledgments

This work was supported by the Wellcome Centre for Mitochondrial Research (203105/Z/16/Z) and AEV was supported by an MRC studentship (MR/K501074/1) and PhD supplementary award as part of the MRC Centre for Neuromuscular Disease (MR/K000608/1) upon initiation of this work and is now in receipt of a Sir Henry Wellcome Fellowship (215888/Z/19/Z). The BBSRC funded the SBFSEM (BB/M012093/1). Work of the authors was supported by NIH grants GM119793, MH122706, and AG066828 to MP.

## Author Contributions

Experimental conception and design: A.E.V., C.L., M.P. Mouse tissue collection: S.K. Methods development and validation: J.F., T.D., K.W., R.L. S.K., C.L., A.K.R, M.P. Acquisition of data: J.F., K.W., T.D. Analysis and interpretation of data: J.F., A.E.V., M.P. Drafting of manuscript: J.F., A.E.V., M.P. Critical revision of manuscript: All authors.

## STAR Methods

### Key Resources Table

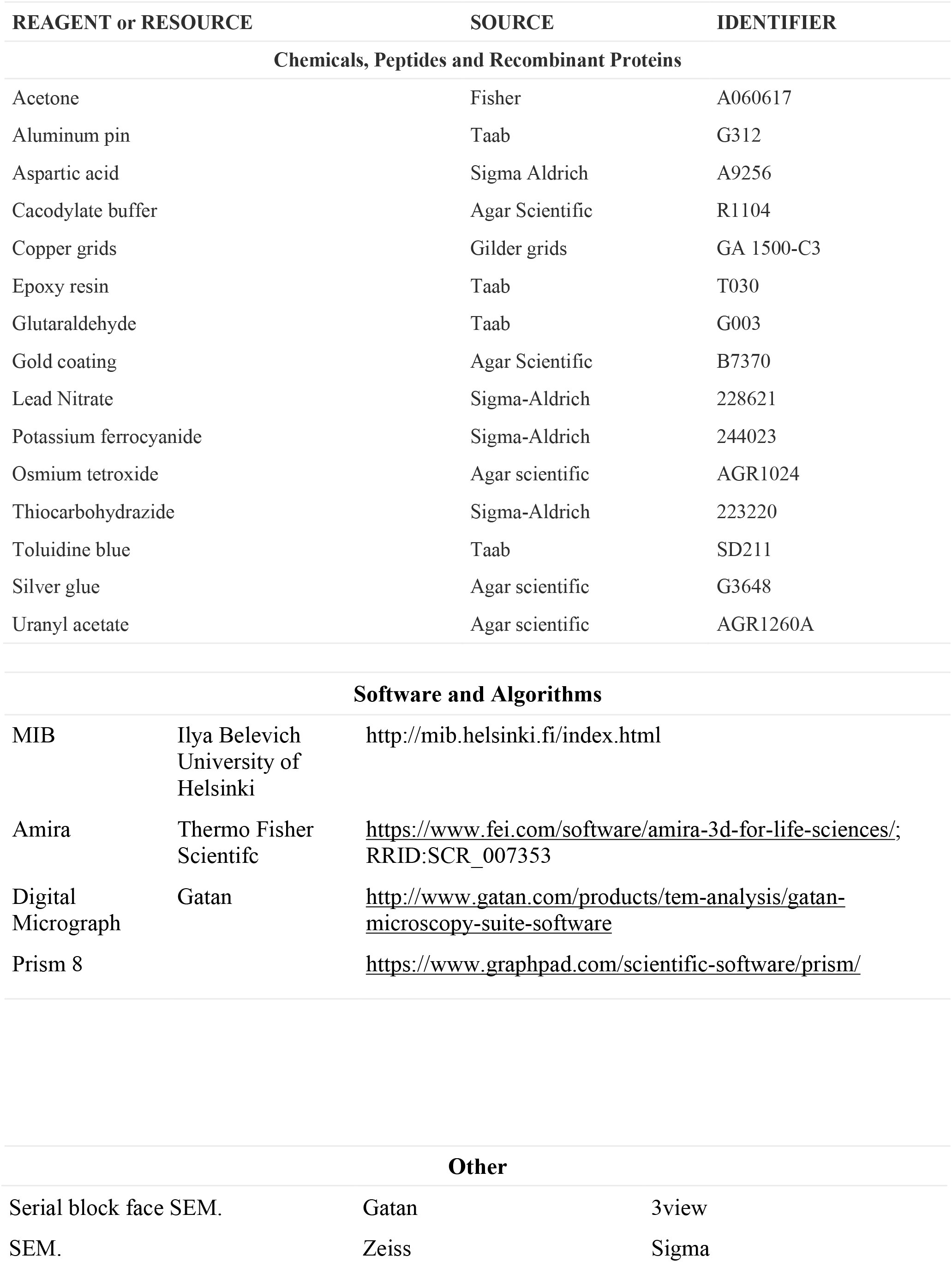

| REAGENT or RESOURCE |  | SOURCE | IDENTIFIER |
| --- | --- | --- | --- |
| Chemicals, Peptides and Recombinant Proteins |  |  |  |
| Acetone |  | Fisher | A060617 |
| Aluminum pin |  | Taab | G312 |
| Aspartic acid |  | Sigma Aldrich | A9256 |
| Cacodylate buffer |  | Agar Scientific | R1104 |
| Copper grids |  | Gilder grids | GA 1500-C3 |
| Epoxy resin |  | Taab | T030 |
| Glutaraldehyde |  | Taab | G003 |
| Gold coating |  | Agar Scientific | B7370 |
| Lead Nitrate |  | Sigma-Aldrich | 228621 |
| Potassium ferrocyanide |  | Sigma-Aldrich | 244023 |
| Osmium tetroxide |  | Agar scientific | AGR1024 |
| Thiocarbohydrazide |  | Sigma-Aldrich | 223220 |
| Toluidine blue |  | Taab | SD211 |
| Silver glue |  | Agar scientific | G3648 |
| Uranyl acetate |  | Agar scientific | AGR1260A |
| Software and Algorithms |  |  |  |
| MIB | Ilya Belevich<br>University of Helsinki | http://mib.helsinki.fi/index.html |  |
| Amira | Thermo Fisher Scientific | https://www.fei.com/software/amira-3d-for-life-sciences/; RRID:SCR_007353 |  |
| Digital Micrograph | Gatan | http://www.gatan.com/products/tem-analysis/gatan-microscopy-suite-software |  |
| Prism 8 |  | https://www.graphpad.com/scientific-software/prism/ |  |
| Other |  |  |  |

|  |  |  |
| --- | --- | --- |
| Serial block face SEM. | Gatan | 3view |
| SEM. | Zeiss | Sigma |

### EXPERIMENTAL MODELS

#### Animals and tissue collection

All experiments involving animals were approved by the Institutional Animal Care and Use Committee of Columbia University Medical Center (IACUC) and were performed in accordance with relevant guidelines and regulations. Mice were maintained under standard conditions, housed in a specific pathogen–free animal house under a 12/12-hour light/dark cycle and were provided food and water ad libitum. All wild type mice C57Bl6 (female) were purchased from Jackson Laboratories (RRID:IMSR_JAX:000664). Young mice were 4 months, (n=3) and old C57BL/6J mice (18 months, n=4) perfused were anesthetized with a mixture of ketamine/xylazine (100/20mg/kg) via intraperitoneal injection.

### METHODS DETAILS

#### Serial block face scanning electron microscopy (SBF-SEM)

Brain thin sections were obtained using a vibratome. The tissues were removed after mice have been perfusion-fixed using buffered 4% glutaraldehyde with 2%PFA in in 0.1M cacodylate buffer and kept in fixative until processing. Then, the samples were processed using a heavy metal protocol adapted from (Deerinck et al., 2010).The samples were washed in 0.1M sodium cacodylate pH7.4 followed by an immersion in 3% potassium ferrocyanide with 2% osmium tetroxide for 1 hour. The samples were put into a contrast enhancer thiocarbohydrazide 0.1% for 20 min, and then 2% osmium tetroxide for 30 min, and finally placed into 1% uranyl acetate overnight at 4 °C. Between each step, all samples were washed in several changes of ddH2O. The day after, the samples were immersed in lead aspartate solution, 0.12 g of lead nitrate in 20 mL aspartic acid, for 30 min at 60 °C. Samples were dehydrated in a graded series of acetone from 25% to 100% and then impregnated in increasing concentrations of Taab 812 hard resin in acetone with several changes of 100% resin. The samples were embedded in 100% fresh resin and left to polymerise at 60 °C for a minimum of 36 hours. After polymerisation, the resin blocks with the tissue were trimmed and the regions of interest (ROIs) were identified and selected by light microscopy first from the thin slices.

Following trimming, samples were placed into a Zeiss Sigma SEM incorporating the Gatan 3view system (Gatan inc., Abingdon, UK) for SBFSEM, which allows sectioning of the block and the collection of serial images in the z-plane. For each mouse, 4 regions of interest (ROIs) in the dentate gyrus (DG, 2 ROIs) and in the cornu-ammonis region 1 (CA1, 2 ROIs) (Figure S1). These ROIs were sectioned and captured in a series of images (400 images per stack) at 50 nm sectioning thickness with dimensions of 4000×2500 pixels and pixel size of 0.01 μm x 0.01 μm. The serial images acquired were handled and processed for segmentation with MIB Microscopy Image Browser, Helsinki (Belevich et al., 2016).

#### 3D-reconstructions and quantitative analysis

Image stacks were exported and analyzed. Somata, nuclei, myelinated axons, dendrites and mitochondria were followed in all three dimensions for reconstructions with MIB.

All the sub-cellular compartments cited above, and mitochondria were manually traced in MIB using the ‘brush’ drawing tool (a manual segmentation tool) over each section and were excluded if they were not completely within the ROI and were analyze. Each material cited above were assigned a colour during the segmentation process. The models have been exported in AMIRA, and for each completely reconstructed structure, the total volume and surface area were extracted and used in analyses. Mitochondrial Complexity Index was calculated using the formula detailed in Vincent *et al*., 2019.

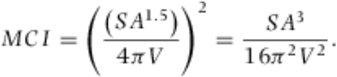

This equation is a three-dimensional equivalent to form factor, to assess mitochondrial morphological complexity.

#### Statistical analyses

Differences in average values of shape descriptions used to assess mitochondrial morphology, effect of aging were analyzed using an Ordinary one–way ANOVA followed by post-hoc tests using the two-stage step up method of Benjamini, Krieger, and Yekutieli to correct for multiple comparisons (p < 0.05, q < 0.05). Differences in average values of shape descriptions used to assess mitochondrial morphology for each single sub-compartment per regions were statistically analysed using nonparametric Mann–Whitney test, because normalisation was not effective. All statistical analyses were performed using the Prism 8 software (GraphPad Software, San Diego, CA, USA).

The experimenter was blinded in all mouse studies.

#### Limitations

Each of the sub-compartment do not correspond to the same cell for each region. The axons of DG and CA1 regions were coming from different places. DG and CA1 molecular layer received inputs from medial-entorhinal cortex (DG) and CA3 (CA1). Furthermore, the representation of each intracellular structures of the cell body is shown in Figure 7A, showing the cytoplasm, the nucleus and all structure with mitochondria in different regions. The volume of each structure was quantified. In order to achieve the segmentation of the full soma without cropping any mitochondria, we often include the start of dendrites and axons in the reconstructions, as it is possible to see in Figure 7A. Also as mentioned previously in the materials and methods, the segmentation was performed to keep the study “blinded”. The characteristic of each animals was unknown, and the selection of each somas was to select ones close to each other. In this way, we show also the differences between all cells segmented (Figure S2).

## Supplemental Table

**Table S1.** Total of Mitochondrial count across neurons sub-cellular compartments in DG and CA1 regions

|  | Axons |  | Dendrites |  | Somas |  |
| --- | --- | --- | --- | --- | --- | --- |
|  | Young | Old | Young | Old | Young | Old |
| DG | 34 | 52 | 55 | 184 | 471 | 777 |
| CA-1 | 48 | 37 | 131 | 146 | 1588 | 1942 |

**Figure S1.**
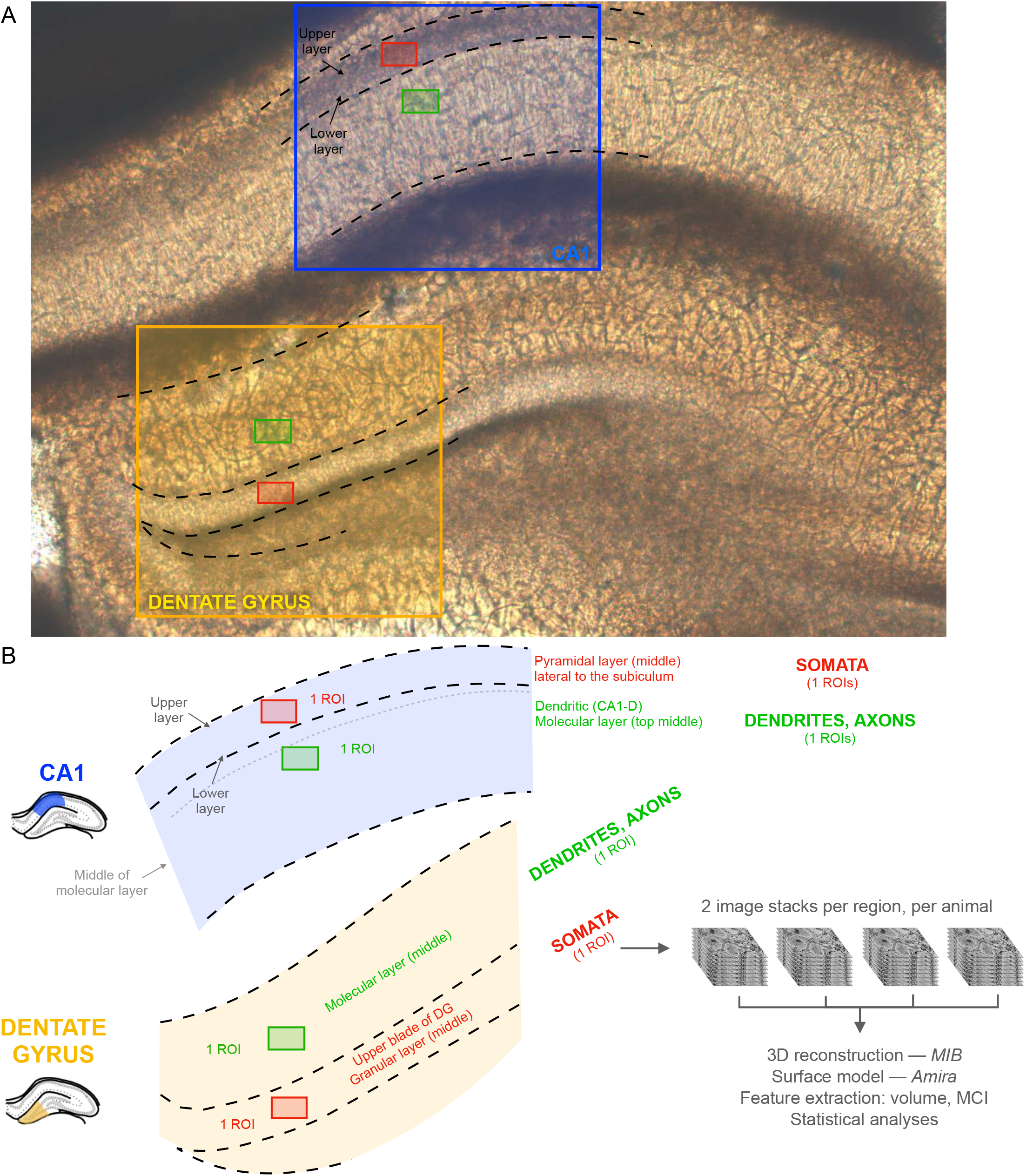
Selection of regions of interest for SBF-SEM of the mouse hippocampus. **(A)** Bright field imaging of the fixed dorsal hippocampus. The DG and CA1 regions uses in analyses are highlighted, and the regions of interest imaged by SBF-SEM are shown as green (molecular layer) and red (granular and pyramidal layers). **(B)** Same regions as in (A) showing anatomical landmarks used for imaging. For each brain region (CA1, DG) and each sub-region (molecular layer, granular/pyramidal layer) of each mouse 2 ROIs were captured, for a total of 4 ROIs per animal. Somata were analysed from ROIs from the granular layer in Dg and from the pyramidal cell layer in CA1. Dendrites and axons were analyzed from the middle of the granular layer of CA near the subiculum and from the upper blade of the DG. Experiments included 7 mice (3 young, 4 old), for a total 28 ROIs and 3D reconstructions from which statistical analyses were performed.

**Figure S2.**
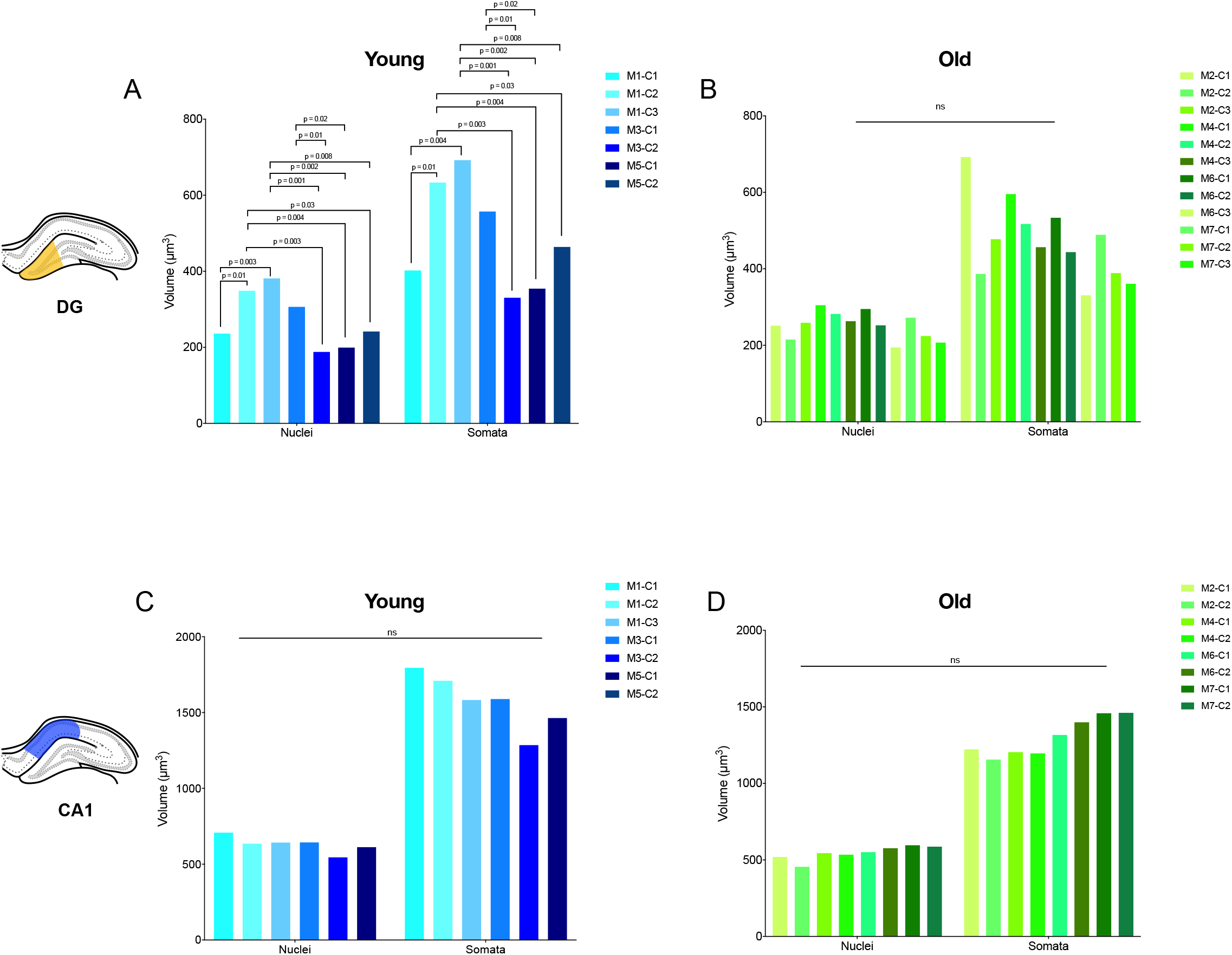
Cell-to-cell differences within the mouse DG and CA1. **(A-B)** Average nuclei and somata volume in individual granule cells in young **(A)** and old **(B)** in the DG. **(C-D)** Average nuclei and somata volume in individual pyramidal cells in young **(C)** and old **(D)** in the CA1. Two–Way ANOVA followed by post-hoc tests using the two-stage step up method of Benjamini, Krieger, and Yekutieli to correct for multiple comparisons (*p* < 0.05, *q* < 0.05).

**Figure S3.**
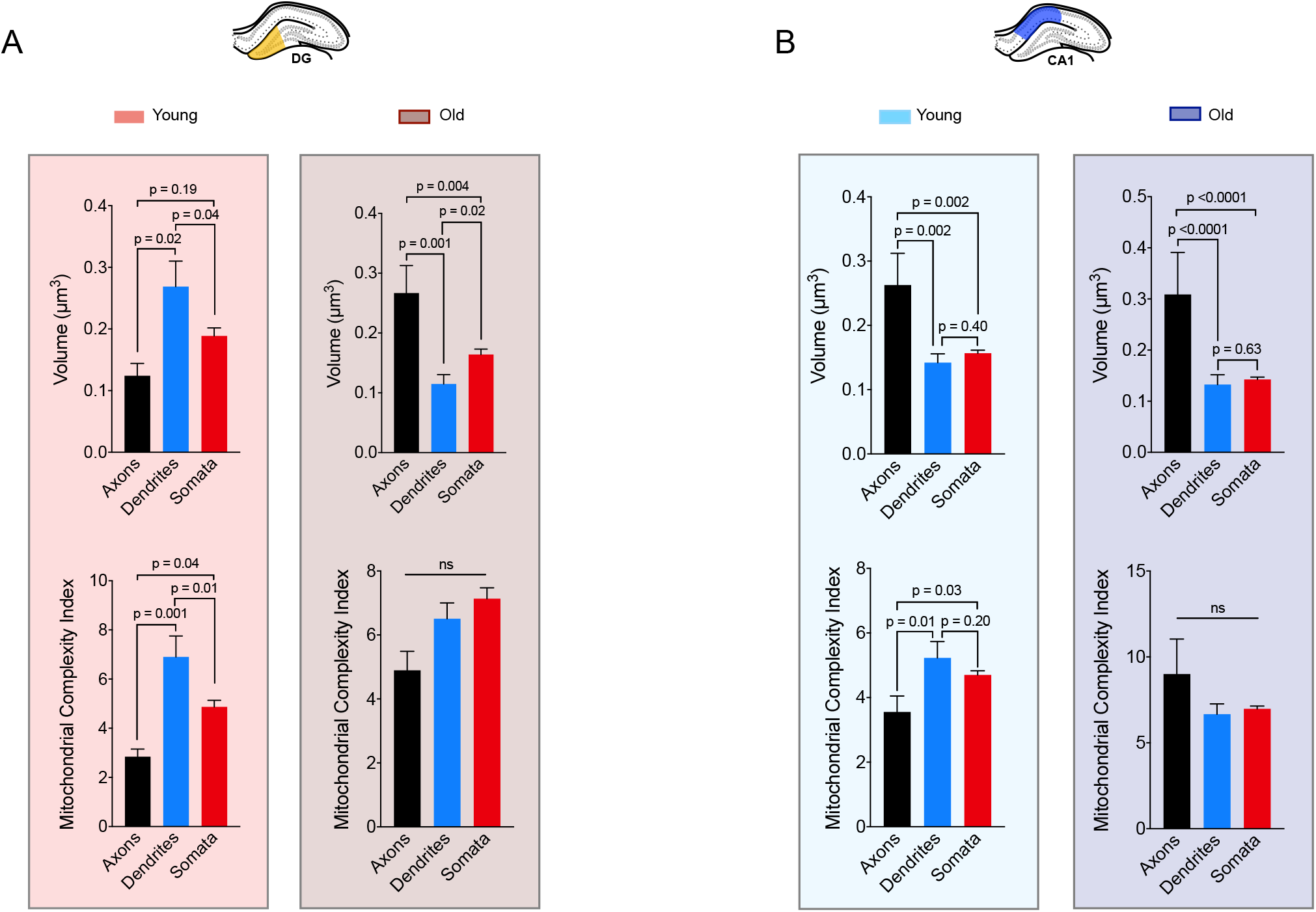
Effect of aging in mitochondrial morphology across sub-cellular compartments. **(A)** Average mitochondrial volume and MCI in DG of axons, dendrites and somata of young and old animals. Left panel Young animals - Right panel Old animals **(B)** Average mitochondrial volume and MCI in CA1 of axons, dendrites and cell bodies of young and old animals. Left panel Young animals - Right panel Old animals Young DG: n= 34 axonal; n=55 dendritic; n=471 somatic mitochondria; Old DG: n= 52 axonal; n=184 dendritic; n=777 somatic. Young CA1: n= 48 axons; n=131 dendrites; n=1588 somatic. Old CA1: n= 37 axonal; n=146 dendritic; n=1930 somatic. n=3 axons, dendrites, and somata analyzed for each region (DG and CA1), in young and old (total 12 of each). Data are means ± SEM. One–Way ANOVA followed by post-hoc tests using the two-stage step up method of Benjamini, Krieger, and Yekutieli to correct for multiple comparisons (*p* < 0.05, *q* < 0.05).

**Figure S4.**
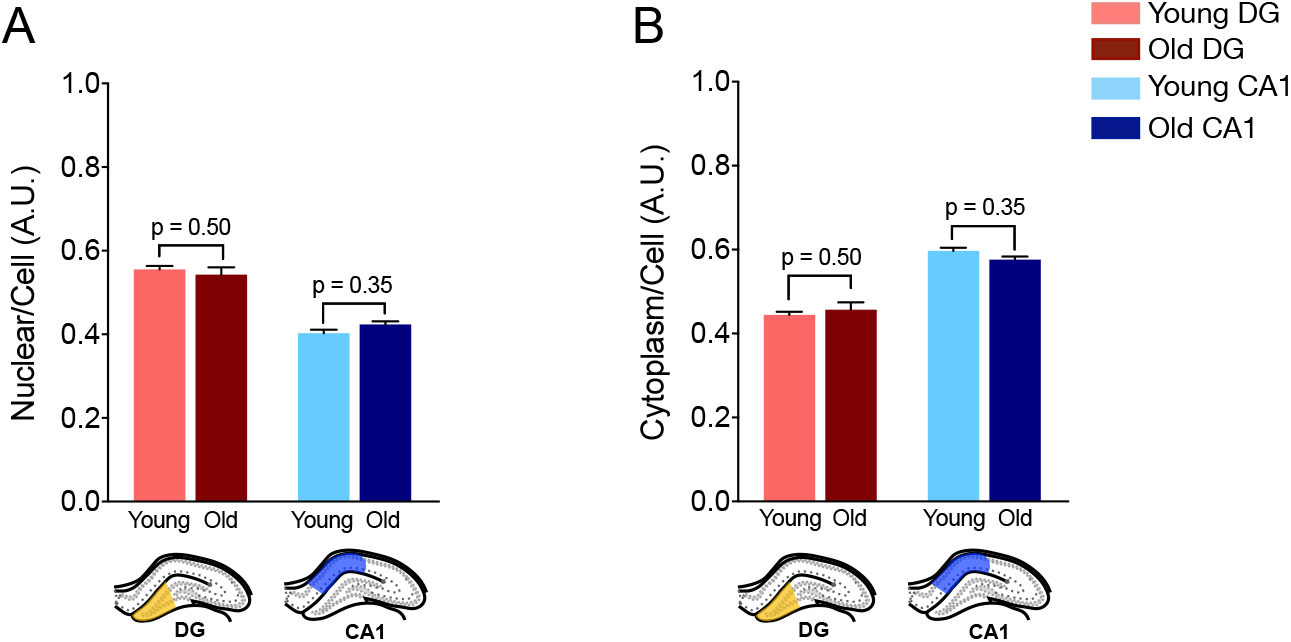
Effect of aging on Nuclear/Cell volume ratio within DG and CA1 region. **(A)** Ratio nuclear/cell and **(B)** ratio cytoplasm/cell Young and Old animals in DG region and CA1 region. One–Way ANOVA followed by post-hoc tests using the two-stage step up method of Benjamini, Krieger, and Yekutieli to correct for multiple comparisons (*p* < 0.05, *q* < 0.05).

